# An osmotic laxative renders mice susceptible to prolonged *Clostridioides difficile* colonization and hinders clearance

**DOI:** 10.1101/2021.07.13.452287

**Authors:** Sarah Tomkovich, Ana Taylor, Jacob King, Joanna Colovas, Lucas Bishop, Kathryn McBride, Sonya Royzenblat, Nicholas A. Lesniak, Ingrid L. Bergin, Patrick D. Schloss

## Abstract

Antibiotics are a major risk factor for *Clostridioides difficile* infections (CDIs) because of their impact on the microbiota. However, non-antibiotic medications such as the ubiquitous osmotic laxative polyethylene glycol (PEG) 3350 also alter the microbiota. Clinicians also hypothesize that PEG helps clear *C. difficile*. But whether PEG impacts CDI susceptibility and clearance is unclear. To examine how PEG impacts susceptibility, we treated C57Bl/6 mice with 5-day and 1-day doses of 15% PEG in the drinking water and then challenged the mice with *C. difficile* 630. We used clindamycin-treated mice as a control because they consistently clear *C. difficile* within 10 days post-challenge. PEG treatment alone was sufficient to render mice susceptible and 5-day PEG-treated mice remained colonized for up to 30 days post-challenge. In contrast, 1-day PEG treated mice were transiently colonized, clearing *C. difficile* within 7 days post-challenge. To examine how PEG treatment impacts clearance, we administered a 1-day PEG treatment to clindamycin-treated, *C. difficile*-challenged mice. Administering PEG to mice after *C. difficile* challenge prolonged colonization up to 30 days post-challenge. When we trained a random forest model with community data from 5 days post-challenge, we were able to predict which mice would exhibit prolonged colonization (AUROC = 0.90). Examining the dynamics of these bacterial populations during the post-challenge period revealed patterns in the relative abundances of *Bacteroides*, *Enterobacteriaceae*, *Porphyromonadaceae*, *Lachnospiraceae*, and *Akkermansia* that were associated with prolonged *C. difficile* colonization in PEG-treated mice. Thus, the osmotic laxative, PEG, rendered mice susceptible to *C. difficile* colonization and hindered clearance.

**Importance:** Diarrheal samples from patients taking laxatives are typically rejected for *Clostridiodes difficile* testing. However, there are similarities between the bacterial communities from people with diarrhea or *C. difficile* infections (CDI) including lower diversity compared to communities from healthy patients. This observation led us to hypothesize that diarrhea may be an indicator of *C. difficile* susceptibility. We explored how osmotic laxatives disrupt the microbiota’s colonization resistance to *C. difficile* by administering a laxative to mice either before or after *C. difficile* challenge. Our findings suggest that osmotic laxatives disrupt colonization resistance to *C. difficile*, and prevent clearance among mice already colonized with *C. difficile*. Considering that most hospitals recommend not performing *C. difficile* testing on patients taking laxatives and laxatives are prescribed prior to administering fecal microbiota transplants via colonoscopy to patients with recurrent CDIs, further studies are needed to evaluate if laxatives impact microbiota colonization resistance in humans.

## Introduction

Antibiotics are a major risk factor for *Clostridioides difficile* infections (CDIs) because they disrupt microbiota colonization resistance (1). However, antibiotics are not the only types of medications that disrupt the microbiota (2–4). Although, other medications (proton pump inhibitors, osmotic laxatives, antimotility agents, and opioids) have been implicated as risk or protective factors for CDIs through epidemiological studies, whether the association is due to their impact on the microbiota is still unclear (5–9).

Many of the non-antibiotic medications associated with CDIs are known to modulate gastrointestinal motility leading to either increased or decreased colonic transit time, which in turn also strongly impacts microbiota composition and function (10, 11). Stool consistency often serves as an approximation of intestinal motility (10). Our group has shown that when *C. difficile* negative samples from patients were separated into two groups based on stool consistency, there were similar microbiota features between samples from CDI patients and *C. difficile* negative patients with diarrhea compared to non-diarrheal samples that were *C. difficile* negative (12). The similar community features between CDI patients and patients with diarrhea included low alpha diversity and only 6 bacterial taxa with higher relative abundances in communities from CDI patients. These results led to the hypothesis that bacterial communities from patients experiencing diarrhea are susceptible to developing CDIs, regardless of how they developed diarrhea.

Depending on the dose administered, osmotic laxatives can lead to diarrhea and temporarily disrupt the human intestinal microbiota (13). The ubiquitous osmotic laxative, polyethylene glycol (PEG) 3350 is found in Miralax, Nulytely, and Golytely and is also commonly used as bowel preparation for colonoscopies. Interestingly, previous studies have shown that treating mice with PEG alone altered microbiota composition, reduced acetate and butyrate production, altered the mucus barrier, and rendered the mice susceptible to *C. difficile* colonization (14–17). The mucus barrier is thought to mediate protection from CDIs by protecting intestinal epithelial cells from the toxins produced by *C. difficile* (18, 19). Whether laxative administration results in more severe CDIs in mice and how long mice remain colonized with *C. difficile* after challenge is unclear.

Beyond susceptibility, PEG is also relevant in the context of treating recurrent CDIs via fecal microbiota transplant (FMT), where a healthy microbiota is administered to the patient to restore colonization resistance. For FMTs that are delivered via colonoscopy, patients typically undergo bowel preparation by taking an osmotic laxative prior to the procedure. Many of the FMT studies to date rationalize the use of laxatives prior to the FMT (20–22) based on a 1996 case study with 2 pediatric patients where the authors suggested in the discussion that the laxative may help flush *C. difficile* spores and toxins from the intestine (23).

Our group has used C57BL/6 mice to characterize how antibiotics disrupt the microbiota and influence *C. difficile* susceptibility and clearance (24–26). Although two groups have now shown that PEG treatment alone renders mice susceptible to *C. difficile* (15, 17), these studies have raised additional questions regarding the dynamics and severity of infection as well as the role of laxative treatment in *C. difficile* clearance. Here, we characterized how long PEG-treated mice remain susceptible, whether PEG treatment results in more sustained *C. difficile* colonization and severe CDI than mice treated with clindamycin, and whether PEG treatment after challenge can promote *C. difficile* clearance. Addressing these questions will better inform how we think about laxatives and diarrhea in the context of CDIs.

## Results

### 5-day laxative treatment led to prolonged *C. difficile* colonization in mice

Building off of previous work that showed treating mice with the osmotic laxative, PEG 3350, rendered mice susceptible to *C. difficile* colonization (15, 17), we decided to test how long *C. difficile* colonization is sustained and how long PEG-treated mice remain susceptible to *C. difficile*. We compared three groups of mice treated with PEG 3350 to one group of mice treated with our standard 10 mg/kg clindamycin treatment, which temporarily renders mice susceptible to *C. difficile* colonization, with mice typically clearing *C. difficile* within 10 days post-challenge (9, 26). All three groups of PEG-treated mice were administered a 15% PEG solution in the drinking water for 5-days. The first group received no additional treatment. The second group was also treated with clindamycin. A third group was allowed to recover for 10 days prior to challenge (Fig. 1A). The PEG treatment resulted in weight loss for the 3 groups of mice, with the greatest change in weight observed on the fifth day of the PEG treatment. The mice recovered most of the lost weight by five days after treatment (Fig. 1B). After either the PEG, clindamycin, or PEG and clindamycin treatment all mice were challenged with 10^5^ *C. difficile* 630 spores (Fig. 1A). All treatments rendered mice susceptible to *C. difficile* colonization. In contrast to the mice that only received clindamycin, PEG-treated mice remained colonized with *C. difficile* at a high level through thirty days post-challenge (Fig. 1C). The clindamycin-treated mice cleared *C. difficile* within ten days post-challenge (Fig. 1C). It was noteworthy that PEG-treated mice were still susceptible to *C. difficile* colonization after a 10-day recovery period, although *C. difficile* was not detectable in most of the group in the initial five days post-challenge (Fig. 1C, S1A). One mouse was found dead on the 6th day post-challenge, presumably due to *C. difficile* as the bacterium became detectable in stool samples from that mouse on the 4th day post-challenge (Fig. S1A, mouse 10). From 8 days post-challenge onward, the density of *C. difficile* stabilized in the 10-day recovery group and remained high through 20-30 days post-challenge (Fig. 1C). Thus, osmotic laxative treatment alone was sufficient to render mice susceptible to prolonged *C. difficile* colonization and PEG-treated mice remained susceptible through ten days post-treatment.

**Figure 1.**
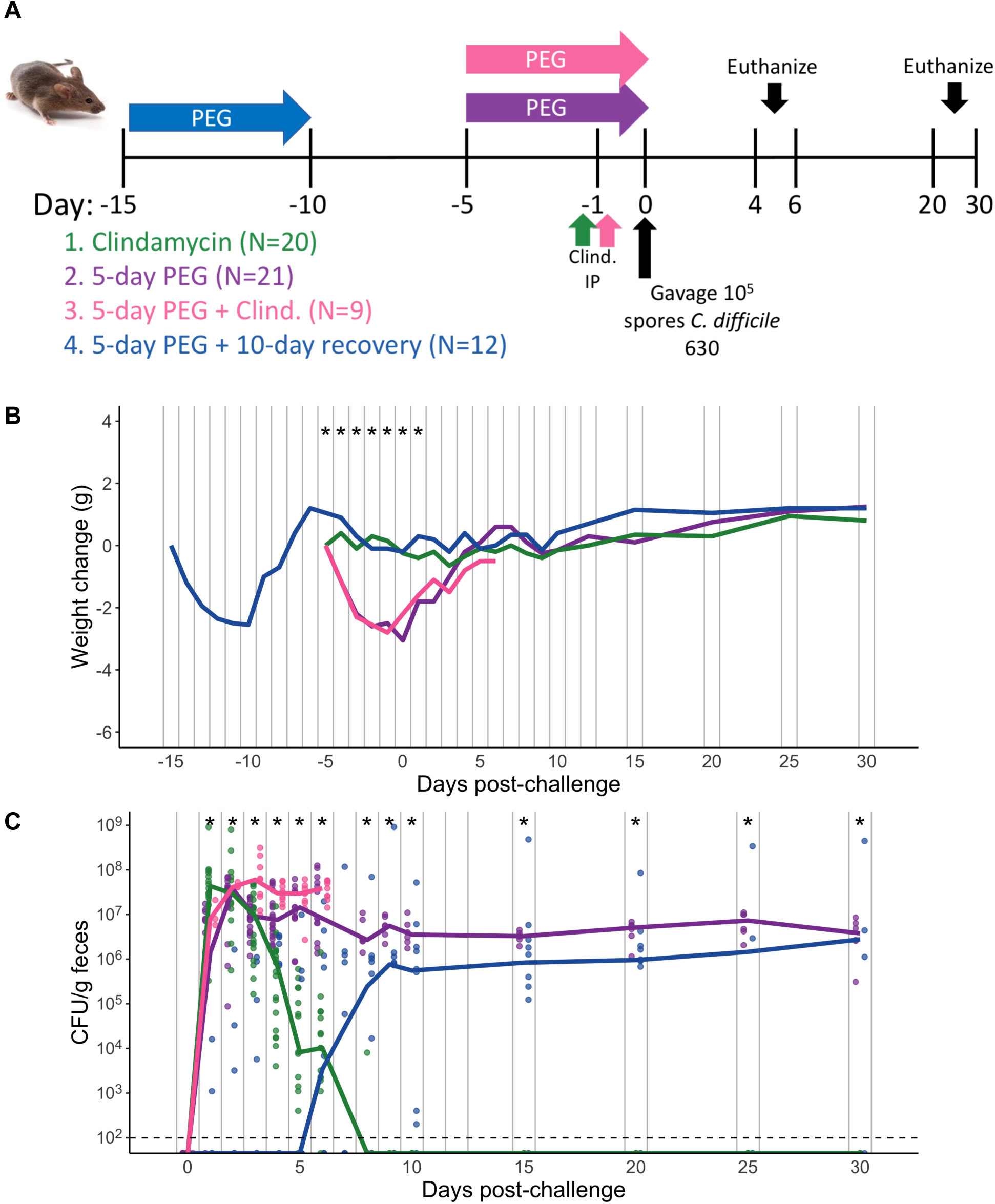
5-day PEG treatment prolongs susceptibility and mice become persistently colonized with *C. difficile*. A. Setup of the experimental time line for experiments with 5-day PEG treated mice consisting of 4 treatment groups. 1. Clindamycin was administered at 10 mg/kg by intraperitoneal injection. 2. 15% PEG 3350 was administered in the drinking water for five days. 3. 5-day PEG plus clindamycin treatment. 4. 5-day PEG plus 10-day recovery treatment. All treatment groups were then challenged with 10^5^ *C. difficile* 630 spores. A subset of mice were euthanized on either 4 or 6 days post-challenge and tissues were collected for histopathology analysis, the remaining mice were followed through 20 or 30 days post-challenge. B. Weight change from baseline weight in groups after treatment with PEG and/or clindamycin, followed by *C. difficile* challenge. C. *C. difficile* CFU/gram stool measured over time via serial dilutions(N = 10-59 mice per time point). The black line represents the limit of detection for the first serial dilution. CFU quantification data was not available for each mouse due to stool sampling difficulties (particularly the day the mice came off of the PEG treatment) or early deaths. For B-C, lines represent the median for each treatment group and circles represent samples from individual mice. Asterisks indicate time points where the weight change or CFU/g was significantly different (*P* < 0.05) between groups by the Kruskal-Wallis test with Benjamini-Hochberg correction for testing multiple time points. The data presented are from a total of 5 separate experiments.

### 5-day laxative treatment differentially disrupted the fecal microbiota compared to clindamycin treatment

Since osmotic laxatives and clindamycin have previously been shown to disrupt the murine microbiota (14–17), we hypothesized the different *C. difficile* colonization dynamics between mice treated with the osmotic laxative or clindamycin were due to the two drugs having differential effects on the microbiota. We profiled the stool microbiota over time by sequencing the V4 region of the 16S rRNA gene to compare changes across treatment groups. We found time (R^2^ = 0.29) and treatment group (R^2^ = 0.21) explained half of the observed variation between fecal communities with most of the remaining variation explained by interactions between treatment group and other experimental variables including time, cage, and sequencing preparation plate (PERMANOVA combined R^2^ = 0.95, *P* < 0.001, Fig. 2A, Data Set S1, sheet 1). None of the treatment groups recovered to their baseline community structure either 10 or 30 days post-challenge, suggesting other community features besides recovery to baseline were responsible for the prolonged *C. difficile* colonization in PEG-treated mice (Fig. 2B).

**Figure 2.**
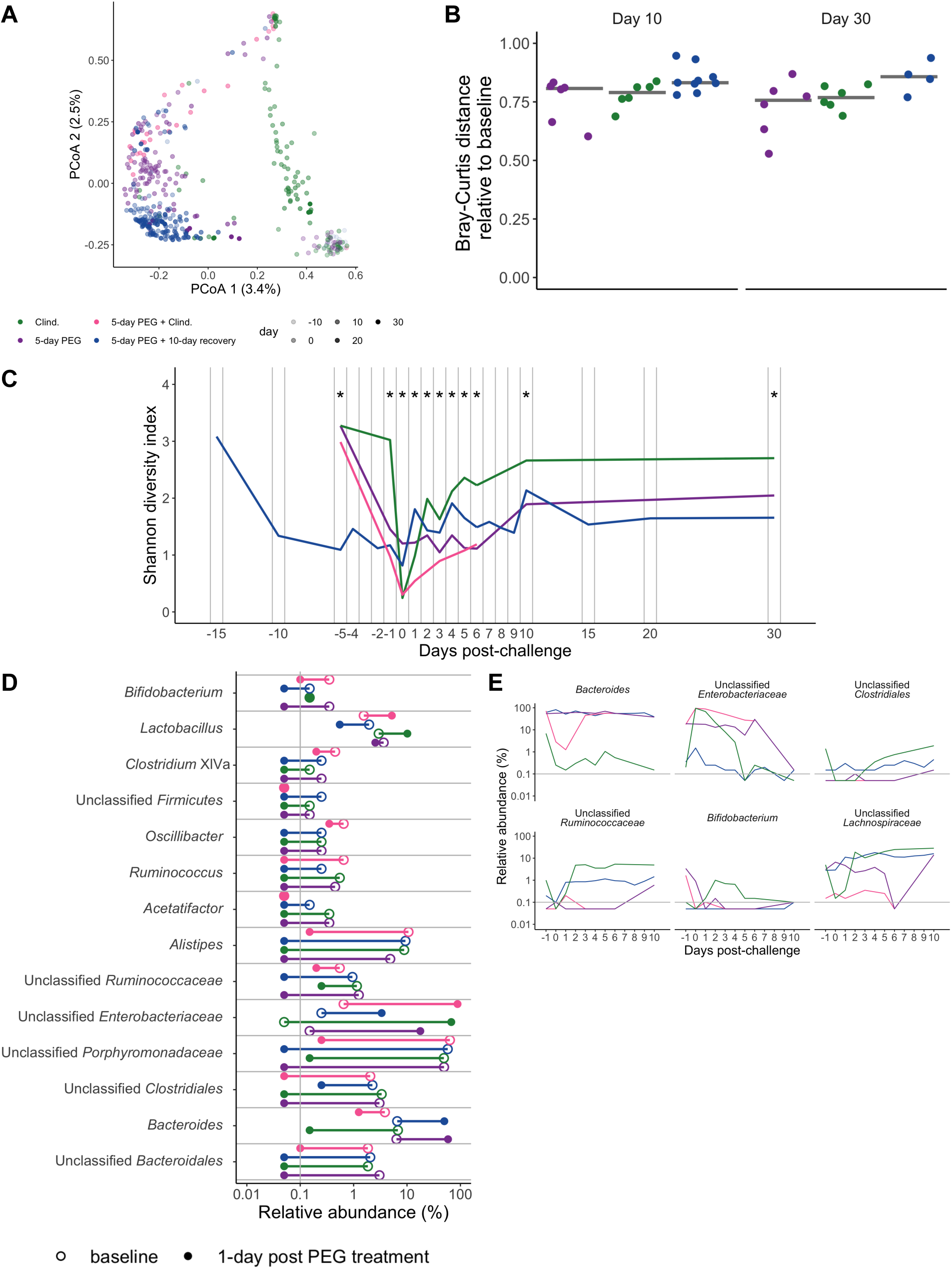
5-day PEG treatment disrupts the stool microbiota for a longer amount of time compared to clindamycin-treated mice. A. Principal Coordinate analysis (PCoA) of Bray-Curtis distances from stool samples collected throughout the experiment. Each circle represents a sample from an individual mouse and the transparency of the symbol corresponds to the day post-challenge. See Data Set S1, sheet 1 for PERMANOVA results. B. Bray-Curtis distances of stool samples collected on either day 10 or 30 post-challenge relative to the baseline sample collected for each mouse (before any drug treatments were administered). The symbols represent samples from individual mice and the line indicates the median value for each treatment group. C. Shannon diversity in stool communities over time. The line indicates the median value for each treatment group (Data Set S1, sheet 2). D. 14 of the 33 genera affected by PEG treatment (Data Set S1, sheet 3). The symbols represent the median relative abundance for a treatment group at either baseline (open circle) or 1-day post treatment (closed circle). Relative abundance data from paired baseline and 1-day post treatment stool sampes from the 5-day PEG and 5-day PEG plus 10-day recovery groups were analyzed by paired Wilcoxan signed-rank test with Benjamini-Hochberg correction for testing all identified genera. The clindamycin and 5-day PEG plus clindamycin treatment groups are shown on the plot for comparison. E. 6 of the 24 genera that were significantly different between the treatment groups over multiple time points (see Data Set S1, sheet 4 for complete list). The 5-day PEG plus clindamycin treatment group was only followed through 6-days post-challenge. Differences between treatment groups were identified by Kruskal-Wallis test with Benjamini-Hochberg correction for testing all identified genera (*, *P* < 0.05). The gray vertical line (D) and horizontal vertical lines (E) indicate the limit of detection.

Because time and treatment group influenced most of the variation between communities, we next explored whether there were differences in community diversity and composition between treatment groups. We examined the alpha diversity dynamics by calculating the communities’ Shannon diversity. Although both clindamycin and PEG treatments decreased diversity, the Shannon diversity was lower in the groups of mice that received PEG treatment compared to those that received clindamycin alone through thirty days post-challenge (Fig. 2C; Data Set S1, sheet 2). We next identified the bacterial genera whose relative abundances shifted after PEG treatment by comparing the baseline samples of mice treated with only PEG to samples from the same mice one day post-PEG-treatment. We found 18 genera whose relative abundances were altered by PEG treatment (Data Set S1, sheet 3). The majority of the bacterial relative abundances decreased after the PEG treatment, but the relative abundance among members of the *Enterobacteriaceae* and *Bacteroides* increased. The increase in *Bacteroides* relative abundance was unique to PEG treated mice, as the *Bacteroides* relative abundance actually decreased in clindamycin treated mice (Fig. 2D). Finally, we identified the genera whose relative abundance differed across treatment groups over multiple time points. Of the 33 genera that were different between treatment groups, 24 genera were different over multiple time points (Fig. 2E, Data Set S1, sheet 4). Thus, PEG had a significant impact on the fecal microbiota that was maintained over time and was distinct from clindamycin treatment.

Because *C. difficile* was not immediately detectable in the stools of the PEG-treated mice that were allowed to recover for 10 days prior to challenge, we decided to examine if there were genera that changed during the post-challenge period. We compared the communities from when *C. difficile* shifted from undetectable at 1 day post-challenge to detectable in the stool samples with the density stabilizing around 8 days post-challenge (Fig. S1A). We found no genera with relative abundances that were significantly different over the two time points (Data Set S1, sheet 5). However, there was also wide variation between individual mice regarding when *C. difficile* became detectable (Fig. S1A) as well as the relative abundances of bacterial genera in the communities (Fig. S1B). For example, two mice had a high relative abundance of *Enterobacteriaceae* throughout the post-challenge period. One mouse died on the sixth day post-challenge and in the other, *C. difficile* was present at a high density from the 4th day post-challenge onward (Fig. S1B). While we did not identify a clear signal to explain the delayed appearance of *C. difficile* in the 5-day PEG mice that were allowed to recover for 10 days prior to challenge, the delay was striking and could reflect changes in microbial activity or metabolites that were not examined in this study.

### 5-day laxative treatment did not promote more severe CDIs despite altering the mucosal microbiota

Given the findings from a previous study that demonstrated PEG treatment disrupts the mucus layer and alters the immune response in mice (16), we decided to examine the impact of PEG treatment on the mucosal microbiota and CDI severity. To evaluate the mucosal microbiota, we sequenced communities associated with tissues collected from the cecum, proximal colon, and distal colon. Similar to what was observed with the stool samples, the alpha diversity was lower in the PEG-treated mice compared to clindamycin treated mice (Fig. 3A, Data Set S1, sheet 6). The alpha diversity of the tissue-associated community increased in PEG-treated mice collected at 20 and 30 days post-challenge (Fig. 3A). Group (R^2^ = 0.33), time point (R^2^ = 0.11), and their interactions with other variables (cage, experiment number, and sample type) explained the majority of the variation observed in mucosal communities (PERMANOVA combined R^2^ = 0.83, *P* < 0.05, Fig. 3B, Data Set S1, sheet 7). We saw the greatest difference in the relative abundance of the mucosal microbiota between treatment groups (clindamycin, 5-day PEG, and 5-day PEG plus clindamycin) at 6 days post-challenge with 10 genera that were significantly different (*P* < 0.05) in all three of the tissue types we collected (cecum, proximal colon, and distal colon; Fig. S2A, Data Set S1, sheet 8, 9, and 10). Interestingly, *Peptostreptococcaceae* (the family with a sequence that matches *C. difficile*) was one of the genera that had a significant difference in relative abundance between treatment groups at 6 days post-challenge. This population was primarily only present in the 5-day PEG treatment group of mice and decreased in the proximal and distal colon tissues over time (Fig. S2B). By 30 days post-challenge, only the relative abundances of *Bacteroides*, *Clostridiales*, *Firmicutes*, and *Ruminococcaceae* were different between treatment groups and only in the cecum tissues (Fig. 3C, Fig. 2E, Data Set S1, sheet 8). Thus, PEG treatment had a significant impact on the mucosal microbiota and we detected *C. difficile* sequences in the cecum, proximal colon, and distal colon tissue communities.

**Figure 3.**
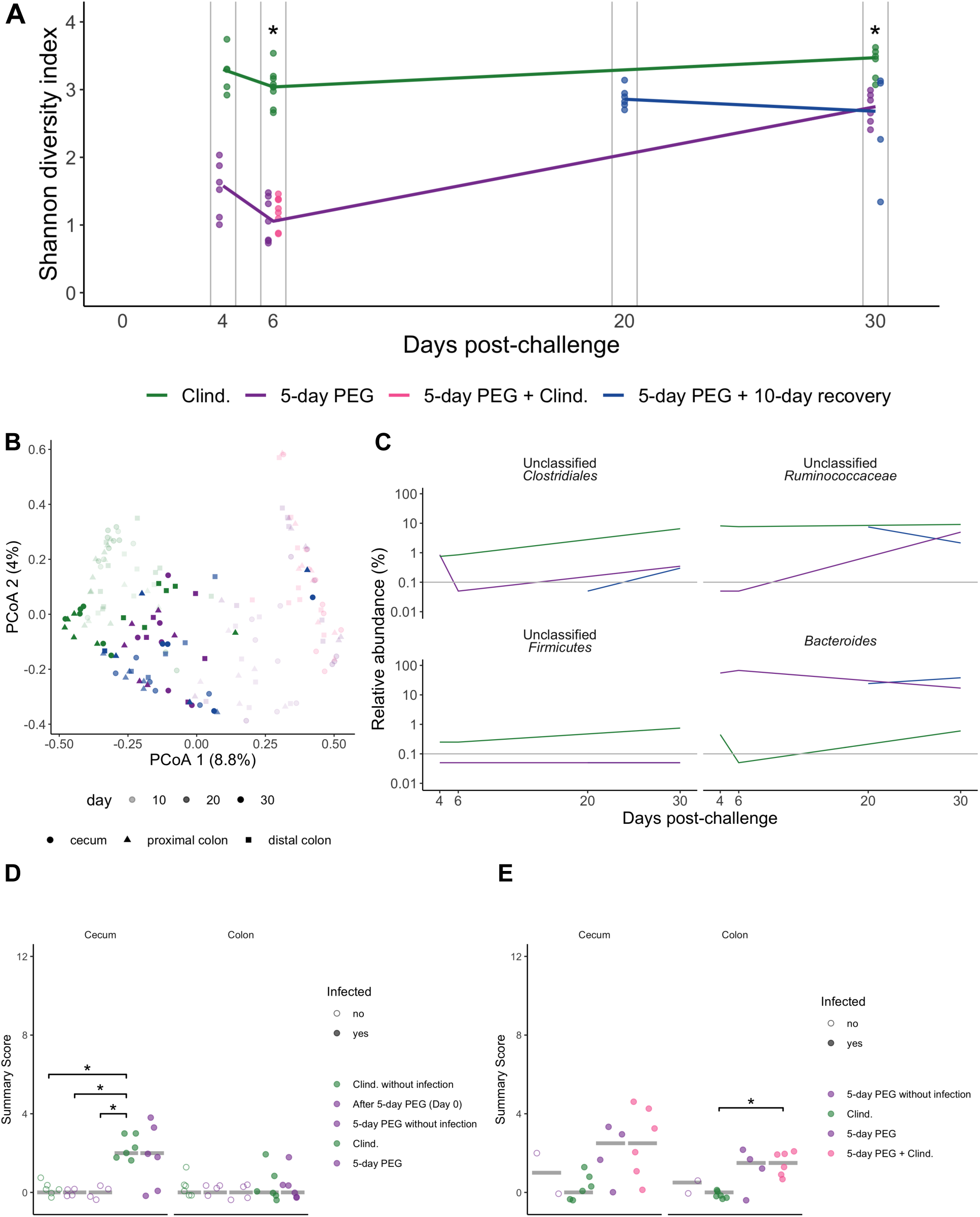
5-day PEG treatment does not result in more severe CDIs, although mucosal microbiota is altered. A. Shannon diversity in cecum communities over time. The colors of the symbols and lines represent individual and median relative abundance values for four treatment groups (Data Set S1, sheet 6). B. PCoA of Bray-Curtis distances from mucosal samples collected throughout the experiment. Circles, triangles, and squares indicate the cecum, proximal colon, and distal colon communities, respectively. Transparency of the symbol corresponds to the day post-challenge that the sample was collected. See Data Set S1, sheet 7 for PERMANOVA results. C. The median relative abundance of the 4 genera that were significantly different between the cecum communities of different treatment groups on day 6 and day 30 post-challenge (Data Set S1, sheet 8). The gray vertical lines indicate the limit of detection. D-E. The histopathology summary scores from cecum and colon H&E stained tissue sections. The summary score is the total score based on evaluation of edema, cellular infiltration, and inflammation in either the cecum or colon tissue. Each category is given a score ranging from 0-4, thus the maximum possible summary score is 12. The tissue for histology was collected at either 4 (D) or 6 (E) days post-challenge with the exception that one set of 5-day PEG treated mock-challenged mice were collected on day 0 post-challenge (first set of open purple circles in D). Histology data were analyzed with the Kruskal-Wallis test followed by pairwise Wilcoxon comparisons with Benjamini-Hochberg correction. *, *P* < 0.05.

**Figure 4.**
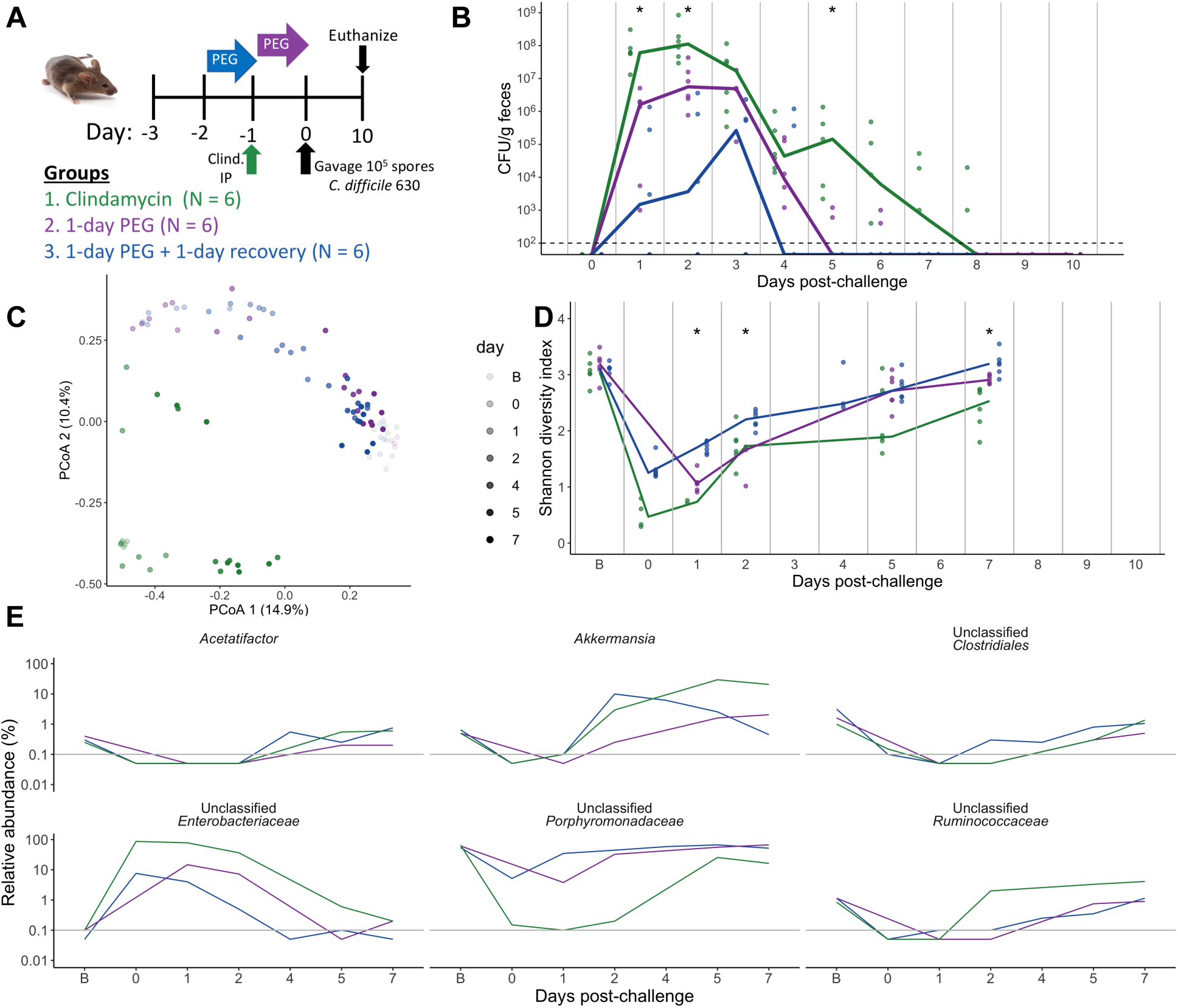
1-day PEG treatment renders mice susceptible to transient *C. difficile* colonization. A. Setup of the experimental time line for the 1-day PEG treated mice consisting of 3 treatment groups. 1. Clindamycin was administered at 10 mg/kg by intraperitoneal injection. 2. 15% PEG 3350 was administered in the drinking water for 1 day. 3. 1-day PEG plus 1-day recovery. The three treatment groups were then challenged with 10^5^ *C. difficile* 630 spores. B. *C. difficile* CFU/gram stool measured over time (N = 12-18 mice per time point) by serial dilutions. The black dashed horizontal line represents the limit of detection for the first serial dilution. For B and D, asterisks indicate time points where there was a significant difference (*P* < 0.05) between treatment groups by Kruskall-Wallis test with Benjamini-Hochberg correction for testing multiple time points. For B-D, each symbol represents a sample from an individual mouse and lines indicate the median value for each treatment group. C. PCoA of Bray-Curtis distances from stool communities collected over time (day: R^2^ = 0.43; group: R^2^ = 0.19, Data Set S1, sheet 11). Symbol transparency represents the day post-challenge of the experiment. For C-E, the B on the day legend or days post-challenge X-axis stands for baseline and represents the sample that was collected prior to any drug treatments. D. Shannon diversity in stool communities over time (Data Set S1, sheet 12). E. Median relative abundances per treatment group for 6 out of the 14 genera that were affected by treatment, but recovered close to baseline levels by 7 days post-challenge (Fig. 3E, Data Set S1, sheets 13 and 14). Paired stool sample relative abundance values either baseline and day 1 or baseline and day 7 were analyzed by paired Wilcoxan signed-rank test with Benjamini-Hochberg correction for testing all identified genera. Only genera that were different between baseline and 1-day post-challenge, but not baseline and 7-days post-challenge are shown. The gray horizontal lines represents the limit of detection.

Because there were differences in the mucosal microbiota, including detectable *C. difficile* sequences in tissues from PEG-treated mice relative to mice treated with clindamycin, we next examined the severity of *C. difficile* challenge by evaluating cecum and colon histopathology (27). However, we found there was no difference in cecum and colon scores between clindamycin and PEG-treated mice that were challenged with *C. difficile* at 4 days post-challenge (Fig. 3D), the time point typically examined in *C. difficile* 630 challenged mice (28). We also looked at 6 days post-challenge because that was when there was a large difference in *C. difficile* density between PEG- and clindamycin-treated mice (Fig. 1C). Although there was a slight difference in the histopathology score of the colon between PEG and clindamycin-treated mice, there was not a signifant difference in the cecum and the overall score was relatively low (1.5 to 2.5 out of 12, Fig. 3E). Therefore, although PEG treatment had a disruptive effect on the mucosal microbiota, the impact of *C. difficile* challenge on the cecum and colon was similar between PEG and clindamycin treated mice.

### *C. difficile* challenge did not have a synergistic disruptive effect on the microbiota of PEG-treated mice

Because *C. difficile* itself can have an impact on the microbiota (29), we also sequenced the tissue and stools of mock-challenged mice treated with clindamycin or PEG. Examining the stools of the mock-challenged mice revealed similar bacterial disruptions as the *C. difficile* challenged mice (Fig. S3A-C). Similarly, there was no difference between the communities found in the tissues of mock and *C. difficile* challenged mice (Fig. S3D-F). Thus, most of the microbiota alterations we observed in the PEG-treated mice were a result of the laxative and not an interaction between the laxative and *C. difficile*.

### 1-day laxative treatment resulted in transient *C. difficile* colonization and minor microbiota disruption

Next, we examined how a shorter osmotic laxative perturbation would impact the microbiome and susceptibility to *C. difficile*. We administered either a 1-day PEG treatment, a 1-day PEG treatment with a 1-day recovery period, or clindamycin to mice before challenging them with *C. difficile* (Fig. 3A). In contrast to the 5-day PEG treated mice, the 1-day PEG groups were only transiently colonized and cleared *C. difficile* by 7 days post-challenge (Fig. 3B). The stool communities of the 1-day PEG treatment groups were also only transiently disrupted, with Shannon diversity recovering by 7 days post-challenge (Fig. 3C-D, Data Set S1, sheets 11 and 12). We found the relative abundances of 14 genera were impacted by treatment, but recovered close to baseline levels by 7 days post-challenge including *Enterobacteriaceae*, *Clostridiales*, *Porpyromonadaceae*, and *Ruminococcaceae* (Fig. 3E, Data Set S1, sheet 13 and 14). These findings suggest the duration of the PEG treatment was relevant, with shorter treatments resulting in a transient loss of *C. difficile* colonization resistance.

### Post-challenge laxative treatment disrupted clearance in clindamycin-treated mice regardless of whether an FMT was also administered

Since a 1-day PEG treatment resulted in a more mild perturbation of the microbiota, we decided to use the 1-day treatment to examine the hypothesis that PEG helps to flush *C. difficile* spores from the intestine. This hypothesis is proposed in the discussion section of FMT studies where bowel prep is part of the preparation undergone by patients receiving FMTs via colonoscopy (20–23). To examine the impact of PEG treatment on *C. difficile* clearance, we treated 4 groups of mice with clindamycin and then challenged all mice with *C. difficile* before administering the following treatments: no additional treatment, 1-day PEG immediately after challenge, and 1-day PEG treatment 3 days after challenge followed by either administration of an FMT or PBS solution by oral gavage (Fig. 5A). Contrary to the hypothesis, all groups of mice that received PEG exhibited prolonged *C. difficile* colonization (Fig. 5B).

**Figure 5.**
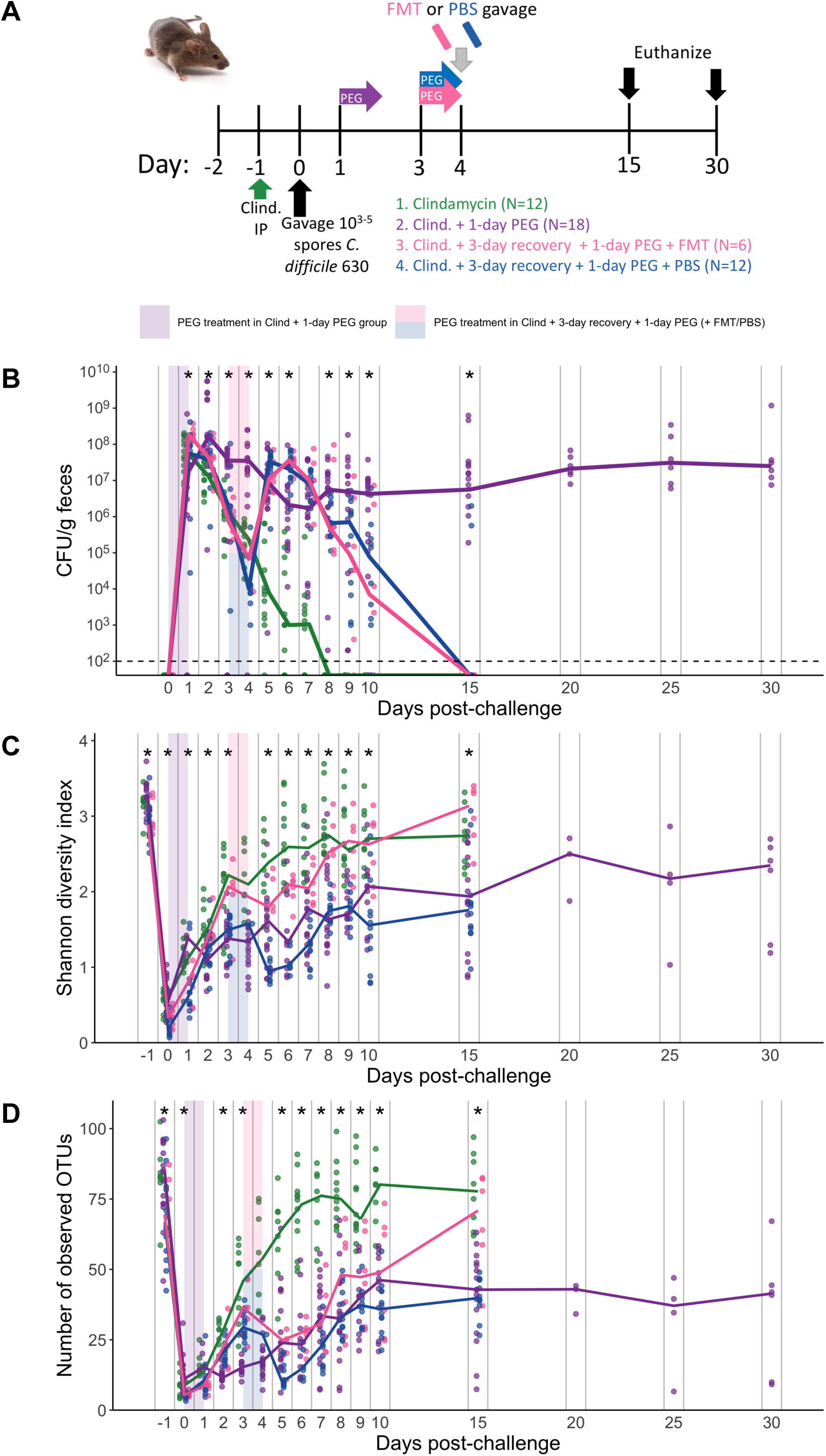
1-day PEG treatment post *C. difficile* challenge prolongs colonization regardless of whether an FMT is also administered. A. Setup of the experimental time line for experiments with post-challenge PEG treated mice. There were a total of 4 different treatment groups. All mice were administered 10 mg/kg clindamycin intraperitoneally (IP) 1 day before challenge with 10^3-5^ *C. difficile* 630 spores. 1. Received no additional treatment (Clindamycin). 2. Immediately after *C. difficile* challenge, mice received 15% PEG 3350 in the drinking water for 1 day. 3-4. 3-days after challenge, mice received 1-day PEG treatment and then received either 100 microliters a fecal microbiota transplant (3) or PBS (4) solution by oral gavage. Mice were followed through 15-30 days post-challenge (only the post-CDI 1-day PEG group was followed through 30 days post-challenge). B. CFU/g of *C. difficile* stool measured over time via serial dilutions. The black line represents the limit of detection for the first serial dilution. C-D. Shannon diversity (C) and richness (D) in stool communities over time (Data Set S1, sheets 15 and 16). B-D. Each symbol represents a stool sample from an individual mouse with the lines representing the median value for each treatment group. Asterisks indicate time points with significant differences (*P* < 0.05) between groups by the Kruskall-Wallis test with a Benjamini-Hochberg correction for testing multiple times points. Colored rectangles indicates the 1-day PEG treatment period for applicable groups. The data presented are from a total of 3 separate experiments.

We were also interested in exploring whether PEG might help with engraftment in the context of FMTs. An FMT was prepared under anaerobic conditions using stool collected from the same group of mice pre-treatment representing the baseline community. The FMT appeared to partially restore Shannon diversity but not richness (Fig. 5C-D, Data Set S1, sheets 15 and 16). Similarly, we saw some overlap between the communities of mice that received FMT and the mice treated with only clindamycin after 5 days post-challenge (Fig. 6A, Data Set S1, sheet 17). The increase in Shannon diversity suggests that the FMT did have an impact on the microbiota, despite seeing prolonged *C. difficile* colonization in the FMT treated mice. However, only the relative abundances of *Bacteroidales* and *Porphyromonadaceae* consistently differed between the mice that received either an FMT or PBS gavage (Fig. 6B). Overall, we found the relative abundances of 24 genera were different between treatment groups over multiple time points (Data Set S1, sheet 18). For example, the relative abundance of *Akkermansia* was increased and the relative abundances of *Ruminococcaceae*, *Clostridiales*, *Lachnospiraceae*, and *Oscillibacter* were decreased in mice that received PEG after *C. difficile* challenge relative to clindamycin treated mice (Fig. 6C). In sum, administering PEG actually prolonged *C. difficile* colonization, including in mice that received an FMT, which only restored 2 bacterial genera.

**Figure 6.**
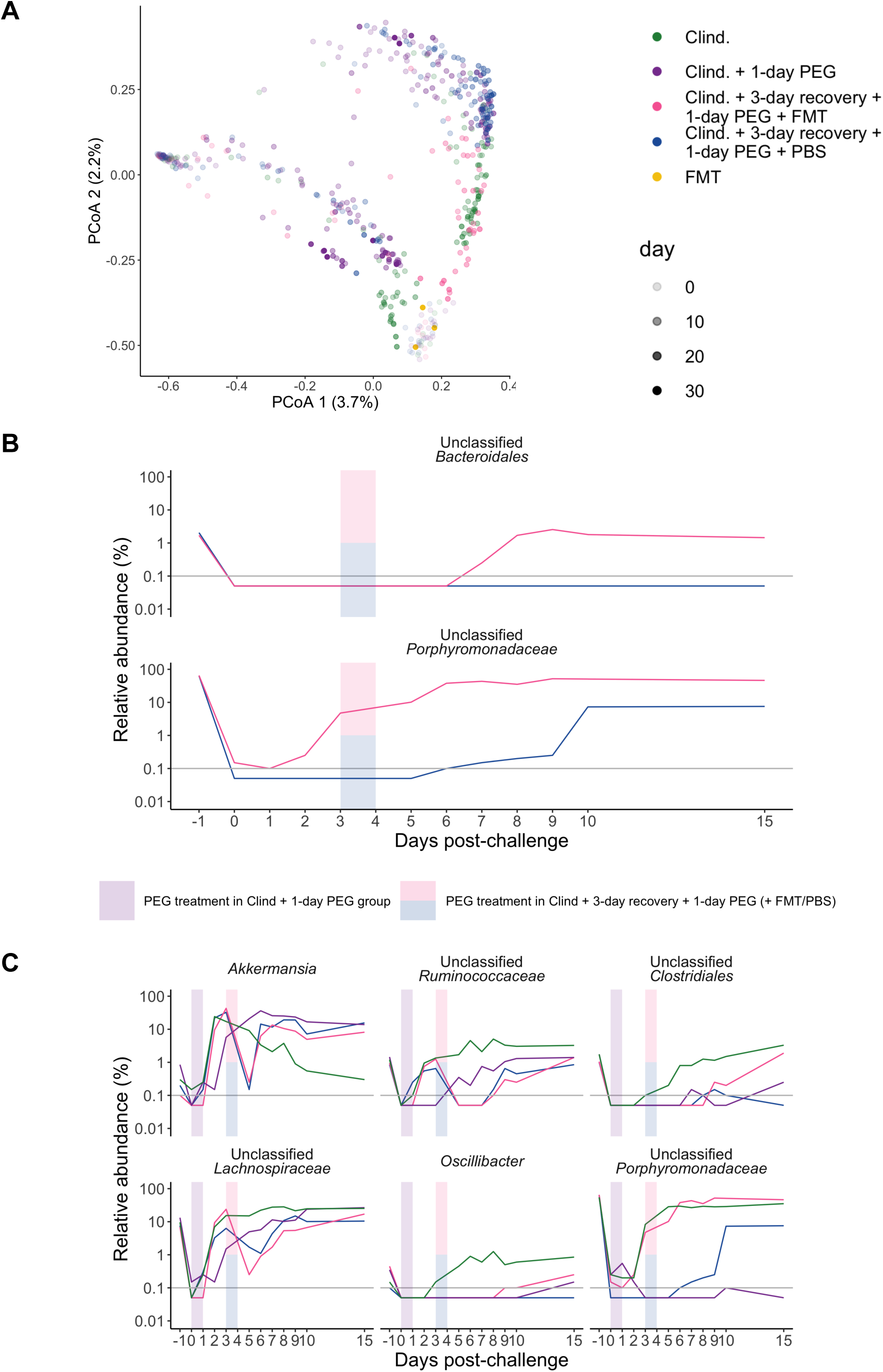
For 1-day PEG treatment post *C. difficile* challenge mice that also receive an FMT only some bacterial genera were restored. A. PCoA of Bray-Curtis distances from stool samples collected over time as well as the FMT solution that was administered to one of the treatment groups. Each circle represents an individual sample, the transparency of the circle corresponds to day post-challenge. See Data Set S1, sheet 17 for PERMANOVA results. B. Median relative abundances of 2 genera that were significantly different over multiple time points in mice that were administered either FMT or PBS solution via gavage C. Median relative abundances of the top 6 out of 24 genera that were significant over multiple time points, plotted over time (see Data Set S1, sheet 18 for complete list). For B-C, colored rectangles indicates the 1-day PEG treatment period for applicable groups. Gray horizontal lines represent the limit of detection. Differences between treatment groups were identified by Kruskal-Wallis test with Benjamini-Hochberg correction for testing all identified genera. For pairwise comparisons of the groups (B), we performed pairwise Wilcoxon comparisons with Benjamini-Hochberg correction for testing all combinations of group pairs.

### Five-day post-challenge community data can predict which mice will have prolonged *C. difficile* colonization

After identifying bacteria associated with the 5-day, 1-day and post-challenge 1-day PEG treatments, we examined the bacteria that influenced prolonged *C. difficile* colonization. We trained 3 machine learning models (random forest, logistic regression, and support vector machine) with bacterial community data from 5 days post-challenge to predict whether the mice were still colonized with *C. difficile* 10 days post-challenge. We chose to predict the status based on communities 5 days post-challenge because that was the earliest time point where we saw a treatment effect in the post-challenge 1-day PEG experiments. The random forest model had the highest performance (median AUROC = 0.90, Data Set S1, sheet 19) and indicated that the 5-day post challenge microbiota was an excellent predictor of prolonged *C. difficile* colonization. Next, we performed a permutation importance test to identify the bacteria that were the top contributors to the random forest model for predicting prolonged *C. difficile* colonization. We selected 10 genera that contributed the most to our model’s performance (Fig. 7A) and examined their relative abundance at 5 days post-challenge, the time point used to predict *C. difficile* colonization status on day 10 (Fig. 7B). Next, we focused on the 5 genera that had a greater than 1% relative abundance in either the cleared or colonized mice and examined how the bacteria changed over time. We found *Enterobacteriaceae* and *Bacteroides* tended to consistently have a higher relative abundance, the relative abundance of *Akkermansia* was initially low and then increased, and *Porphyromonadaceae* and *Lachnospiraceae* had a lower relative abundance in the mice with prolonged colonization compared to the mice that cleared *C. difficile* (Fig. 7C). Together these results suggest a combination of low and high abundance bacterial genera influence the prolonged colonization observed in 5-day PEG and post-challenge 1-day PEG treated mice.

**Figure 7.**
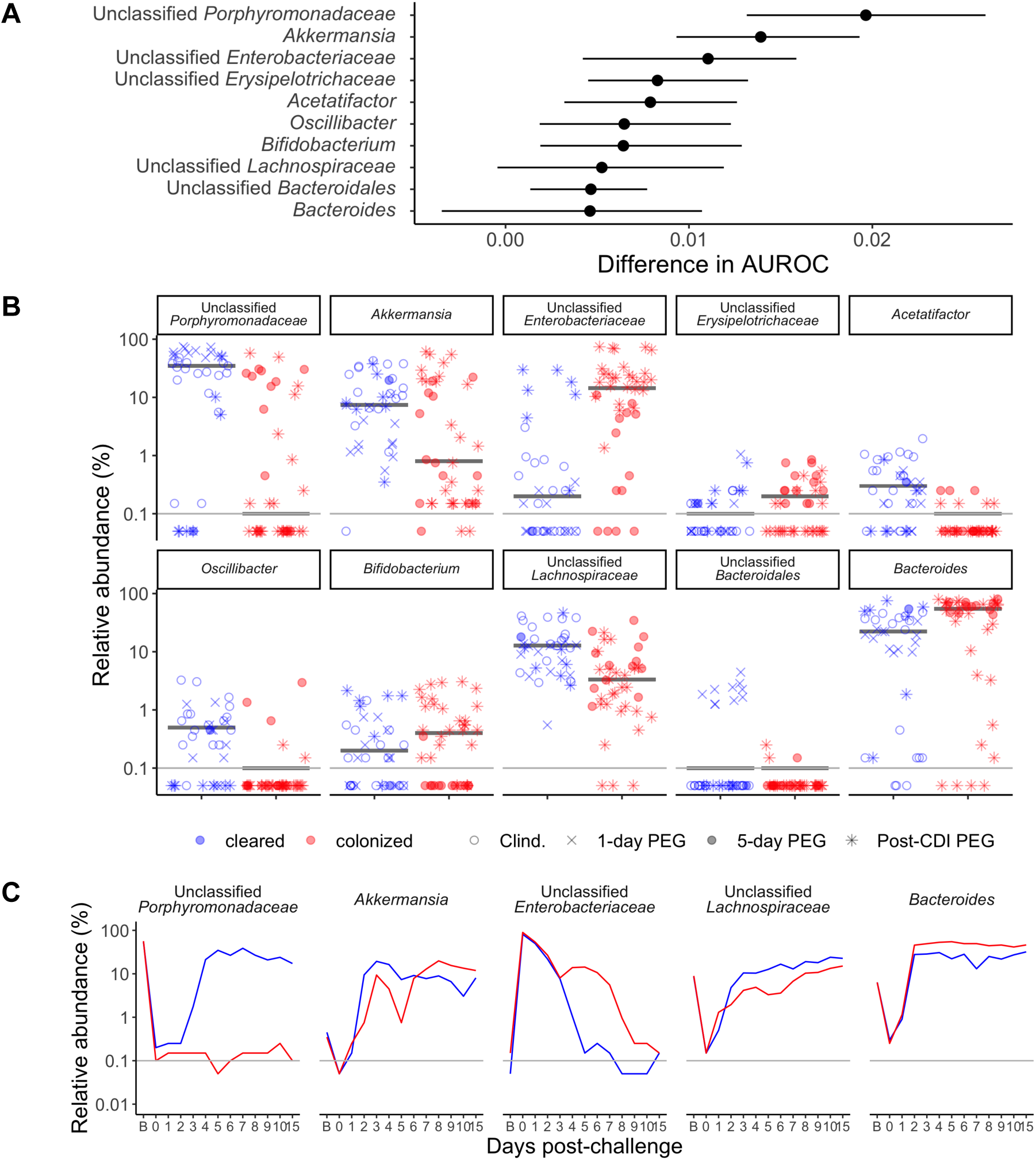
Specific microbiota features associated with prolonged *C. difficile* colonization in PEG treated mice. A. Top ten bacteria that contributed to the random forest model trained on 5-day post-challenge community relative abundance data, predicting whether mice would still be colonized with *C. difficile* 10 days post-challenge. The median (point) and interquartile range (lines) change in AUROC when the bacteria were left out of the model by permutation feature importance analysis. B. The median relative abundances of the top ten bacteria that contributed to the random forest classification model at 5 days post-challenge . Red indicates the mice were still colonized with *C. difficile* while blue indicates mice that cleared *C. difficile* 10 days post-challenge and the black horizontal line represents the median relative abundance for the two categories. Each symbol represents a stool sample from an individual mouse and the shape of the symbol indicates whether the PEG-treated mice received a 5-day (Fig. 1-3), 1-day (Fig. 4) or post-challenge PEG (Fig. 5-6) treatment. C. The median relative abundances of the 5 genera with greater than 1% median relative abundance in the stool community over time. For B-C, the gray horizontal lines represents the limit of detection.

## Discussion

While the disruptive effect of antibiotics on *C. difficile* colonization resistance is well established, the extent to which other drugs such as laxatives disrupt colonization resistance was unclear. By following mice treated with an osmotic laxative over time, we found that a 5-day PEG treatment before challenge resulted in prolonged *C. difficile* colonization, while a 1-day PEG treatment resulted in transient colonization without the use of antibiotics. The differences in *C. difficile* colonization dynamics between the 5- and 1-day PEG treated mice were associated with differences in the degree to which treatments disrupted the microbiota. Additionally, the intestinal communities of 5-day PEG treated mice did not regain colonization resistance after a 10-day recovery period. In contrast to the other 5-day PEG treatment groups, *C. difficile* was not immediately detectable in the stools of most of the mice in the 10-day recovery group. We also examined the impact of PEG treatment after *C. difficile* challenge. In opposition to the hypothesis suggested by the literature, we found that PEG treatment prolonged colonization relative to mice that only recieved clindamycin treatment. We identified patterns in the relative abundances of *Bacteroides*, *Enterobacteriaceae*, *Akkermansia*, *Porphyromonadaceae*, and *Lachnospiraceae* that were associated with prolonged *C. difficile* colonization (Fig. 8). Overall, our results demonstrated that osmotic laxative treatment alone rendered mice susceptible to *C. difficile* colonization and the duration of colonization depended on the length of PEG treatment and whether treatment was administered before or after challenge.

**Figure 8.**
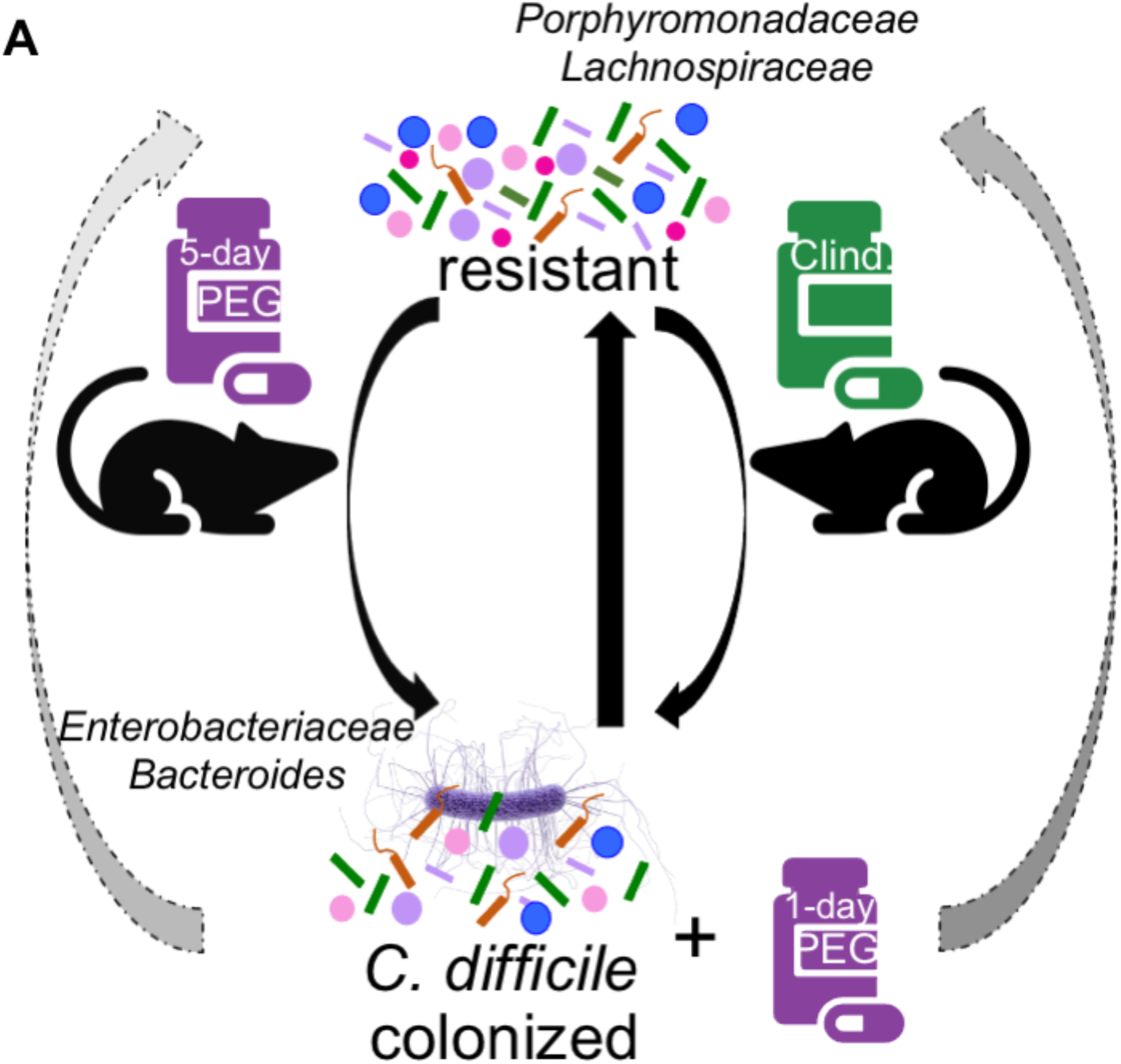
Schematic summarizing findings. The gut microbiota of our C57Bl/6 mice is resistant to *C. difficle* but treatment with either the antibiotic, clindamycin, or the osmotic laxative, PEG 3350, renders the mice susceptible to *C. difficile* colonization. Recovery of colonization resistance in clindamycin-treated mice is relatively straightforward and the mice clear *C.difficile* within 10 days post-challenge. However, for mice that received either a 5-day PEG treatment prior to *C. difficile* challenge or a 1-day PEG treatment post-challenge recovery of colonization resistance was delayed because most mice were still colonized with *C. difficile* 10 days post-challenge. We found increased relative abundances of *Porphyromonadaceae* and *Lachnospiraceae* were associated with recovery of colonization resistance, while increased relative abundances of *Enterobacteriaceae* and *Bacteroides* were associated with prolonged *C. difficle* colonization.

In addition to altering composition, laxative treatment may alter microbiota-produced metabolites. A previous study demonstrated that a 5-day treatment of 10% PEG depleted acetate and butyrate and increased succinate compared to untreated mice (15). While we did not perform metabolomic analysis, we did see bacteria known to produce beneficial metabolites were depleted in mice that cleared *C. difficile* compared to mice with prolonged colonization. For example, *Oscillibacter valericigenes* can produce the SCFA valerate (30), and separate studies demonstrated valerate is depleted after clindamycin treatment and inhibited *C. difficile* growth *in vitro* and in C57BL/6 mice (31, 32). Similarly, *Acetatifactor* can produce acetate and butyrate (33), SCFAs that are decreased in mice with prolonged *C. difficile* infection after antibiotic treatment (34). Thus protective bacteria and their metabolites could be depleted by osmotive laxative treatment depending on the timing and duration of treatment.

One possible explanation for the prolonged *C. difficile* colonization in 5-day PEG treated mice, might be due to the bacteria’s persistence in the mucosal compartment. In fact, it has been hypothesized that *C. difficile* biofilms may serve as reservoirs for recurrent infections (35) and *C. difficile* biofilms in the mucus layer were recently identified in patients as aggregates with *Fusobacterium nucleatum* (36). There was an interesting pattern of increased *Enterobacteriaceae*, *Bacteroides*, and *C. difficile* in both the stool and mucosal communities of PEG-treated mice suggesting a potential synergy. *Bacteroides* has the potential to degrade mucus and the osmotic laxative may have allowed *Bacteroides* to colonize the mucosal niche by degrading mucin glycans with glycosyl hydrolases that are absent in *C. difficile* (37). *Bacteroides* persistence in the mucosal tissue might also have helped *Enterobacteriaceae* to colonize the region, as a synergy between mucus-degrading *B. fragilis* and *E. coli* has previously been described (38). A separate study demonstrated *C. difficile* was present in the outer mucus layer and associated with *Enterobacteriaceae* and *Bacteroidaceae* using fluorescent in situ hybridization (FISH) staining (39). However, protective roles for *Bacteroides* have also been demonstrated. For example, *B. fragilis* prevented CDI morbidity in a mouse model and inhibited *C. difficile* adherence *in vitro* (40). In coculture experiments, *B. longum* decreased *C. difficile* biofilm formation while *B. thetaiotamicron* enhanced biofilm formation (41) and *B. dorei* reduced *C. difficile* growth in a 9-species community *in vitro* (42). Therefore, whether *Bacteroides* is detrimental or beneficial in the context of *C. difficile* infection or colonization is still unclear, but the niche and interactions with other bacteria may contribute.

*Akkermansia* is also a mucin degrader with potentially beneficial or detrimental roles depending on context in other diseases (43, 44). In our study, the relative abundance of *Akkermansia* shifted over time between groups of mice that either cleared *C. difficile* or had prolonged colonization. In the stool it was initially increased in mice that cleared *C. difficile*, but shifted after 5-days post-challenge so that it was increased in mice that had prolonged colonization. In the context of CDIs, some studies suggest a protective role (45, 46), while others suggest a detrimental role because *Akkermansia* was positively correlated with *C. difficile* (47–50). Because the relative abundance of *Akkermansia* was dynamic in our study, it is unclear whether *Akkermansia* helps with clearance of *C. difficile* or allows it to persist. A better understanding how *C. difficile* interacts with the mucosal microbiota may lead to insights into CDIs, asymptomatic *C. difficile* carriage, and colonizatiion resistance.

Despite identifying an altered compositional profile that included higher relative abundance of the *C. difficile* sequence in the mucosal tissues of mice treated with 5-day PEG compared to the clindamycin group, we did not see a difference in histopathology scores between the groups. One reason there was no difference could be the *C. difficile* strain used, *C. difficile* 630 results in mild histopathology summary scores in mice compared to VPI 10463 despite both strains producing toxin in mice (51). Part of our hypothesis for why there could have been increased histopathology scores in PEG-treated mice was because PEG was previously shown to disrupt the mucus layer in mice. However, recent studies demonstrated that broad spectrum antibiotics can also disrupt the host mucosal barrier in mice (52, 53). Further research is needed to tease out the interplay between medications that influence the mucus layer and different strains of *C. difficile* in the context of CDIs.

It is more difficult to interpret what are findings mean in the context of *C. difficile* colonization resistance in human patients. The main difficulty being that most hospitals recommend not performing *C. difficile* testing if the patient is currently taking a laxative. This recommendation is in accordance with the Infectious Diseases Society of America and Society for Healthcare Epidemiology of America guidelines (54). The rationale behind the recommendation is that patients taking laxatives may be asymptomatically colonized with *C. difficile*, resulting in unnecessary antibiotic treatment (55–57). Furthermore, some studies identified laxatives as a risk factor for developing CDIs or recurrent CDIs (58–60) and a recent study found the proportion of severe CDIs was similar between patients taking and not taking laxatives (61). However, there have also been some studies that suggest laxatives are not a risk factor for developing CDIs (62, 63). Although, it is unclear whether laxatives impact CDI susceptibility in human paitents, it is clear that laxatives also have a transient impact on the human microbiota (13, 64–67). Additional studies to examine the relationship between laxatives, *C. difficile* colonization, and CDIs are warranted.

Considering laxatives are also used to prepare patients when administering fecal microbiota transplants via colonoscopy to treat recurrent CDIs, it will be important to determine whether osmotic laxatives impact *C. difficile* clearance in the human intestinal tract. It is still unclear what the best administration route is because there have been no studies designed to evaluate the best administration route for FMTs (68). Nevertheless, results from the FMT National Registry where 85% of FMTs were delivered by colonoscopy demonstrate FMTs are highly effective treatment for recurrent CDIs with 90% achieving resolution in the 1 month follow-up window (69). A surprising number of studies continue to hypothesize that PEG or bowel preparation can clear *C. difficile* spores and toxins despite the paucity of supporting evidence (20–23). There was even a clinical trial (NCT01630096) designed to examine whether administering PEG 3350 (NuLYTELY) prior to antibiotic treatment reduced disease severity that started recruitment in 2012 (70), but no results have been posted to date. Here we sought to evaluate the impact of treating *C. difficile* colonized mice with PEG (with or without FMT) and found clearance was delayed. Further studies are needed to understand the impact of osmotic laxatives on *C. difficile* colonization resistance and clearance in human patients receiving FMTs.

We have demonstrated that osmotic laxative treatment alone has a substantial impact on the microbiota and rendered mice susceptible to prolonged *C. difficile* colonization in contrast to clindamycin-treated mice. The duration and timing of the laxative treatment impacted the duration of *C. difficile* colonization, with only 5-day PEG and post-challenge 1-day PEG treatments prolonging colonization compared to clindamycin treated mice. Further studies are warranted to ascertain whether laxatives have a similar impact on *C. difficile* colonization resistance on the human microbiota.

## Acknowledgements

We thank members of the Schloss lab for feedback on planning the experiments and data presentation. We thank Andrew Henry for help with media preparation and bacterial culture and the Microbiology and Immunology department’s postdoctoral association writing group members for their feedback on manuscript drafts. We also thank the Unit for Laboratory Animal Medicine at the University of Michigan for maintaining our mouse colony and providing the institutional support for our mouse experiments. Finally, we thank Kwi Kim, Austin Campbell, and Kimberly Vendrov for their help in maintaining the Schloss lab’s anaerobic chamber. This work was supported by the National Institutes of Health (U01AI124255). ST was supported by the Michigan Institute for Clinical and Health Research Postdoctoral Translation Scholars Program (UL1TR002240 from the National Center for Advancing Translational Sciences).

## Materials and Methods

### Animals

All experiments were approved by the University of Michigan Animal Care and Use Committee IACUC (protocol numbers PRO00006983 and PRO00008975). All mice were C57Bl/6 and part of the Schloss lab colony which was established in 2010 with mice donated from Vincent Young’s lab colony (established with mice purchased from The Jackson Laboratory in 2002). We used 7-19 week old female mice for all experiments. This allowed us to break up littermates and distribute them as evenly as possible across treatment groups in order to minimize microbiota differences prior to starting treatments with medications. During the experiment, mice were housed at a density of 2-3 mice per cage, with the majority of cages limited to two mice.

### Drug treatments

For PEG treament groups, fifteen percent PEG 3350 (Miralax) was administered in the drinking water for either 5 or 1-day periods depending on the experiment. PEG solution was prepared fresh every 2 days in distilled water and administered to the mice in water bottles. Clindamycin treatment groups received distilled water in water bottles during the PEG-treatment periods, with the water being changed at the same frequency. For clindamycin treatment, groups of mice received 10 mg/kg clindamycin (Sigma-Aldrich) via intraperitoneal injection. All PEG treatment groups received a sham intraperitoneal injection containing filter sterilized saline.

### *C. difficile* challenge model

Mice were challenged with 25 microliters containing 10^5^ *C. difficile* 630 spores, except for 1 experiment where the concentration was 10^3^ (Fig. 5A). All mock challenged mice received 25 ul vehicle solution (Ultrapure water). A Dymax stepper pipette was used to administer the same challenge dose to mice via oral gavage. Mice were weighed daily throughout the experiment and stool was collected for quantifying *C. difficile* CFU and 16S rRNA gene sequencing. There were two groups of mice that received either a PBS or fecal microbiota transplant (FMT) gavage post-PEG treatment. The fecal microbiota transplant was prepared with stool samples collected from the mice in the experiment prior to the start of any treatments. The stool samples were transferred to an anaerobic chamber and diluted 1:10 in reduced PBS and glycerol was added to make a 15% glycerol solution. The solution was then aliquoted into tubes and stored at -80°C until the day of the gavage. An aliquot of both the FMT and PBS solutions were also set aside in the -80°C for 16S rRNA gene sequencing. The day of the gavage, aliquots were thawed and centrifuged at 7500 RPM for 1 minute. The supernatant was then transferred to a separate tube to prevent the gavage needle from clogging with debris during gavage. The PBS solution that was administered to the other group was also 15% glycerol. Each mouse was administered 100 microliters of either the FMT or PBS solution via gavage. When we refer to mice that cleared *C. difficile*, we mean that no *C. difficile* was detected in the first serial dilution (limit of detection: 100 CFU). In some experiments, we collected tissues for 16SrRNA gene sequencing, histopathology, or both. For 16S rRNA gene sequencing, we collected small snips of cecum, proximal colon, and distal colon tissues in microcentrifuge tubes, snap froze in liquid nitrogen, and stored at -80°C. For histopathology, cecum and colon tissues were placed into separate cassettes, fixed, and then submitted to McClinchey Histology Labs (Stockbridge, MI) for processing, embedding, and hematoxylin and eosin (H&E) staining.

### *C. difficile* quantification

Stool samples from mice were transferred to an anaerobic chamber and serially diluted in reduced PBS. Serial dilutions were plated onto taurocholate-cycloserine-cefoxitin-fructose agar (TCCFA) plates plates and counted after 24 hours of incubation at 37°C. Stool samples collected from the mice on day 0 post-challenge were also plated onto TCCFA plates to ensure mice were not already colonized with *C. difficile* prior to challenge.

### 16S rRNA gene sequencing

Stool samples were stored at -80°C and were placed into 96-well plates for DNA extractions and library preparation. DNA was extracted using the DNeasy Powersoil HTP 96 kit (Qiagen) and an EpMotion 5075 automated pipetting system (Eppendorf). For library preparation, each plate had a mock community control (ZymoBIOMICS microbial community DNA standards) and a negative control (water). The V4 region of the 16S rRNA gene was amplified with the AccuPrime Pfx DNA polymerase (Thermo Fisher Scientific) using custom barcoded primers, as previously described (71). The PCR amplicons were normalized (SequalPrep normalizatin plate kit from Thermo Fisher Scientific), pooled and quantified (KAPA library quantification kit from KAPA Biosystems), and sequenced with the MiSeq system (Illumina).

### 16S rRNA gene sequence analysis

All sequences were processed with mothur (v. 1.43) using a previously published protocol (71, 72). Paired sequencing reads were combined and aligned with the SILVA (v. 132) reference database (73) and taxonomy was assigned with a modified version of the Ribosomal Database Project reference sequences (v. 16) (74). The error rate for are sequencing data was 0.0559% based on the 17 mock communities we ran with the samples. Samples were rarefied to 1,000 sequences, 1,000 times for alpha and beta diversity analyses in order to account for uneven sequencing across samples. All but 3 of the 17 water controls had fewer than 1000 sequences. PCoAs were generated based on Bray-Curtis Index distance matrices. Permutational multivariate analysis of variance (PERMANOVA) tests were performed on mothur-generated Bray-Curtis distance matrices with the adonis function from the vegan R package (75).

### Histopathology

H&E stained sections of cecum and colon tissues collected at either 0, 4, or 6 days post-challenge were coded to be scored in a blinded manner by a board-certified veterinary pathologist (ILB). Slides were evaluated using a scoring system developed for mouse models of *C. difficile* infection (51). Each slide was evaluated for edema, cellular infiltration, and inflammation and given a score ranging from 0-4. The summary score was calculated by combining the scores from the 3 categories and ranged from 0-12.

### Classification model training and evaluation

We used the mikropml package to train and evaluate models to predict *C. difficile* colonization status 10 days post-challenge where mice were categorized as either cleared or colonized (76, 77). We removed the *C. difficile* genus relative abundance data prior to training the model. Input community relative abundance data at the genus level from 5 days post-challenge was used to generate random forest, logistic regression, and support vector machine classification models to predict *C. difficile* colonization status 10 days post-challenge. To accommodate the small number of samples in our data set we used 50% training and 50% testing splits with repeated 2-fold cross-validation of the training data for hyperparamter tuning. Permutation importance was performed as described previously (78) using mikropml (76, 77) with the random forest model because it had the highest AUROC value.

### Statistical analysis

R (v. 4.0.2) and the tidyverse package (v. 1.3.0) were used for statistical analysis (79, 80). Kruskal-Wallis tests with Bejamini-Hochberg correction for testing multiple time points were used to analyze differences in *C. difficile* CFU, mouse weight change, and alpha diversity between treatment groups. Paired Wilcoxon rank signed rank tests were used to identify genera impacted by treatments on matched pairs of samples from 2 time points. Bacterial relative abundances that varied between treatment groups at the genus level were identified with the Kruskal-Wallis test with Benjamini-Hochberg correction for testing all identified OTUs, followed by pairwise Wilcoxon comparisons with Benjamini-Hocherg correction.

### Code availability

Code for data analysis and generating this paper with accompanying figures is available at https://github.com/SchlossLab/Tomkovich_PEG3350_mSphere_2021.

### Data availability

The 16S rRNA sequencing data have been deposited in the National Center for Biotechnology Information Sequence Read Archive (BioProject Accession no. PRJNA727293).

---

**Figure S1.**
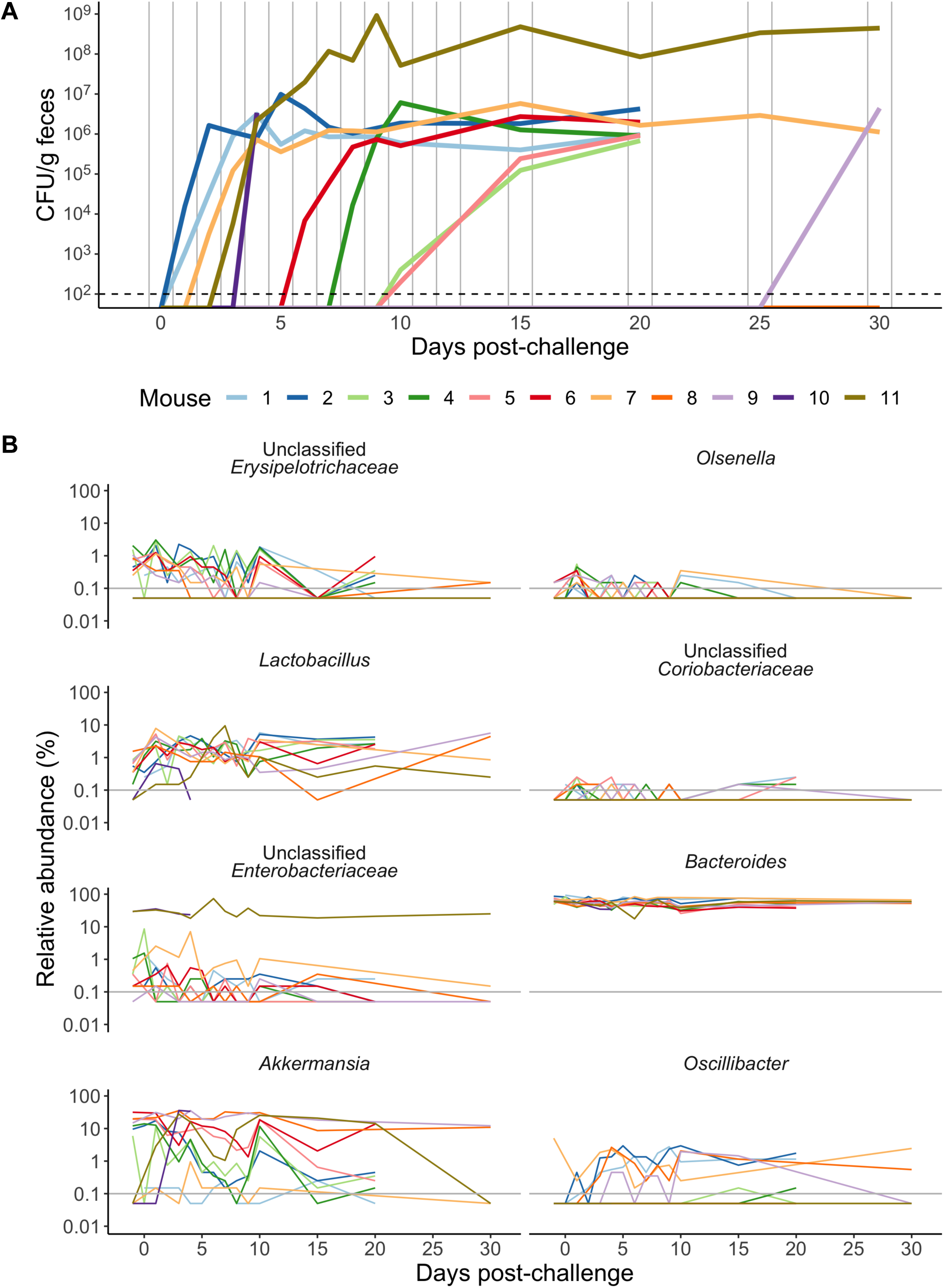
Microbiota dynamics post-challenge in the 5-day PEG treatment plus 10-day recovery mice. A. *C. difficile* CFU/g over time in the stool samples collected from 5-day PEG treated mice that were allowed to recover for 10 days prior to challenge. Same data presented in Fig. 1C, but the data for the other 3 treatment groups have been removed and each line represents the CFU over time for an individual mouse. Mouse 10 was found dead 6 days post-challenge. B. Relative abundances of eight bacterial genera from day 0 post-challenge onward in each of the 10-day recovery mice. We analyzed samples from day 0 and day 8 post-challenge, which represented the time points where mice were challenged with *C. difficile* and when the median relative *C. difficile* CFU stabilized for the group using the paired Wilcoxan signed-rank test, but no genera were significantly different after Benjamini-Hochberg correction (Data Set S1, sheet 5).

**Figure S2.**
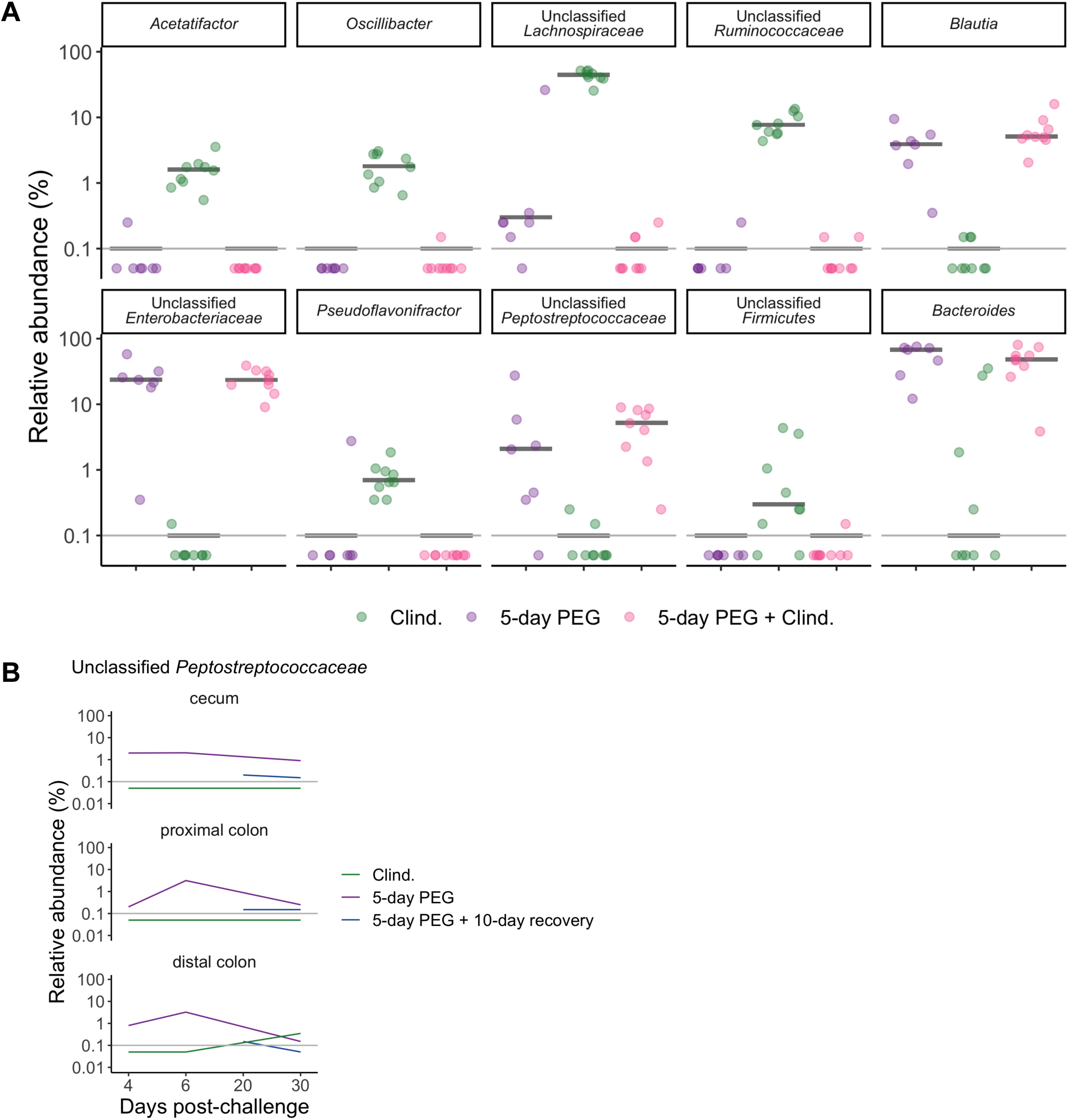
PEG treatment still has a large impact on the mucosal microbiota 6 days post-challenge. A. The relative abundances of the 10 bacterial genera that were significantly different between treatment groups at 6 days post-infection in the cecum tissue (the relative abundances of the 10 genera were also significantly different in the proximal and distal colon tissues, Data Set S1, sheets 8, 9, and 10). Each symbol represents a tissue sample from an individual mouse, the black horizontal lines represents the median relative abundances for each treatment group. B. The relative abundance of *Peptostreptococacceae* in the three types of tissue sample communities over time. For A-B, the gray horizontal lines represent the limit of detection.

**Figure S3.**
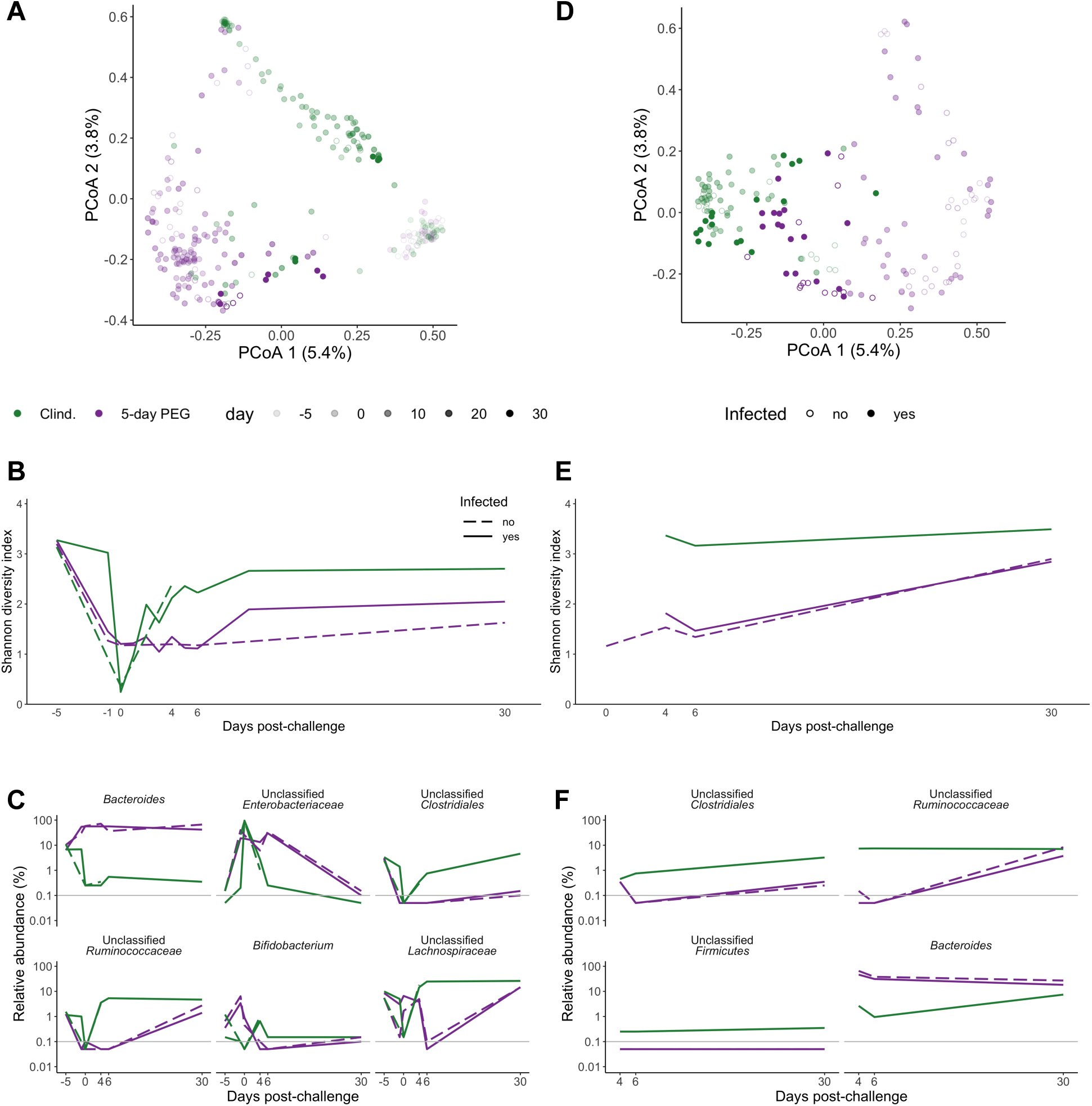
*C. difficile* challenge does not enhance the disruptive effect of PEG on the microbiota. A, D. PCoAs of the Bray-Curtis distances from the stool (A) and tissue (D) communities from mock- and *C. difficile*-challenged treatment groups. Each symbol represents a sample from an individual mouse with open and closed circles representing mock and *C. difficile*-challenged mice, respectively. B, E. Median Shannon diversity in stool (B) and tissue (E) communities collected over time. C, F. The median relative abundances of genera that were significantly different between the *C. difficile* challenged treatment groups in either the stool (Fig. 2E) or cecum tissue (Fig. 3C) communities in the stool (C) and tissue (F) communities from mock- and *C. difficile*-challenged mice. For B-F, the dashed and solid lines represent the median value for mock and *C. difficile*-challenged mice, respectively. For E-F, tissues from mock-challenged clindamycin treated mice were only collected 4 days post-challenge so there is no dashed line for this group.

## Data Set S1

**Data Set S1, Sheets 1-19. Excel workbook with 19 sheets.**

**Data Set S1, Sheet 1. PERMANOVA results for the stool communities from mice in the 5-day PEG subset.**

**Data Set S1, Sheet 2. Shannon diversity analysis for the stool communities from mice in the 5-day PEG subset.**

**Data Set S1, Sheet 3. Genera with relative abundances impacted by PEG treatment based on stool communities of 5-day PEG treated mice.**

**Data Set S1, Sheet 4. Genera with relative abundances that vary between treatment groups in the stool communities from mice in the 5-day PEG subset.**

**Data Set S1, Sheet 5. Statistical analysis results for genera with relative abundances that varied in stool communities in the 5-day PEG plus 10-day recovery mice between the day 1 and day 8 time points.**

**Data Set S1, Sheet 6. Shannon diversity analysis for the cecum communities from mice in the 5-day PEG experiments.**

**Data Set S1, Sheet 7. PERMANOVA results for the tissue communities from mice in the 5-day PEG subset.**

**Data Set S1, Sheet 8. Genera with relative abundances that vary between treatment groups in the cecum communities from mice in the 5-day PEG esubset.**

**Data Set S1, Sheet 9. Genera with relative abundances that vary between treatment groups in the proximal colon communities from mice in the 5-day PEG subset.**

**Data Set S1, Sheet 10. Genera with relative abundances that vary between treatment groups in the distal colon communities from mice in the set of 5-day PEG subset.**

**Data Set S1, Sheet 11. PERMANOVA results for the stool communities from mice in the set of 1-day PEG subset.**

**Data Set S1, Sheet 12. Shannon diversity analysis for the stool communities from mice in the 1-day PEG experiments.**

**Data Set S1, Sheet 13. Genera with different relative abundances between the baseline and day 1 time points in the 1-day PEG subset.**

**Data Set S1, Sheet 14. Genera with different relative abundances between the baseline and day 7 time points in the 1-day PEG subset..**

**Data Set S1, Sheet 15. Shannon diversity analysis for the stool communities from mice in the post-challenge PEG experiments.**

**Data Set S1, Sheet 16. Richness analysis for the stool communities from mice in the post-challenge PEG experiments.**

**Data Set S1, Sheet 17. PERMANOVA results for the stool communities from mice in the post-challenge PEG subset.**

**Data Set S1, Sheet 18. Genera with relative abundances that vary between treatment groups in the stool communities from mice in the post-challenge PEG subset.**

**Data Set S1, Sheet 19. AUROC results for the 100 different seeds from each of the 3 models tested.**

